# High pathogenicity island, *aer* and *sit* operons: a “ménage à trois” in *Escherichia coli* extra-intestinal virulence

**DOI:** 10.1101/2022.11.25.517969

**Authors:** Guilhem Royer, Olivier Clermont, Bénédicte Condamine, Sara Dion, Marco Galardini, Erick Denamur

**Author notes:** Corresponding author: Erick Denamur.

## Abstract

The intrinsic virulence of extra-intestinal pathogenic *Escherichia coli* is attributed to numerous chromosome and/or plasmid-borne virulence associated genes (VAGs), encoding diverse functions as adhesins, toxins, protectins and iron capture systems, which occur in specific genetic backgrounds. Little is however known on their respective contribution to virulence. Here, by analyzing genomes of 232 sequence type complex (STc) 58 strains, we show that virulence quantified in a mouse model of sepsis emerged in a sub-group of STc58 due to the presence of the siderophore encoding high-pathogenicity island (HPI). When extending our analysis to 370 *Escherichia* strains we show that full virulence is associated with the presence of the *aer* or *sit* operons, in addition to the HPI. The prevalence of these operons, their co-occurrence and genomic location depend on the strain phylogeny. Selection of lineage-dependent specific associations of VAGs argues for strong epistatic interactions shaping the emergence of virulence in *E. coli*.

## Introduction

*Escherichia coli* extra-intestinal infections represent a considerable burden both in human and veterinary medicines ^1,2^. *E. coli* is the first bacterial pathogen in humans responsible for deaths associated with antibiotic resistance ^3^. The population structure of *E. coli* is globally clonal^4^ with the delineation of several phylogenetic groups and numerous sequence types^5^. The virulence of the strains is mainly due to the presence of virulence associated genes (VAGs) present on the chromosome and/or plasmids and coding for adhesins, protectins, iron capture systems, invasins and toxins. As *E. coli* acts as an opportunistic extra-intestinal pathogen^6^, the severity of the disease is mainly due to host characteristics as the age, the presence of comorbidities and to the portal of entry for bloodstream infections^7,8^. In this context, intrinsic extra-intestinal virulence of the strains is often assessed in animal models. The chicken systemic infection model via air sacs^9^ and the mouse sepsis assay^10,11^ are robust, reproducible and widely used models.

Using the mouse model, it has been shown that the effects of these VAGs on virulence are cumulative^12^ and dependent on the genetic background of the strain^13,14^, an argument for epistatic interaction between loci across the genome. Extra-intestinal pathogenic *E. coli* (ExPEC) with multiple VAGs belong mainly to phylogroups B2 and D and are highly virulent in the mouse model^10^. Recently, using this assay coupled to a genome wide association study (GWAS) in a collection of 370 strains representative of the genetic diversity of the *Escherichia* genus, the iron capture systems were shown to have a major role in virulence with the high-pathogenicity island (HPI) having the greatest association, followed by the aerobactin (*iuc/iut* genes) and *sit* operons^15^. Interestingly, the aerobactin siderophore system as well as the Sit iron transport system can be encoded by Col-like plasmid genes and/or chromosomal genes^16^ whereas the HPI is a chromosomal genomic island which encodes for a siderophore (yersiniabactin) mediated iron-uptake system^17^. Col-like plasmids are modular plasmids encompassing numerous VAGs, including also salmochelin (*iro* genes), another siderophore.

Although most of the extra-intestinal pathogenic clones belong to the phylogroups B2, D and to a lesser extend C and F^6^, a lineage responsible for human extra-intestinal infections and antibiotic resistance belonging to the phylogroup B1 has recently emerged, namely the ST58^18^ according to the Warwick University nomenclature, belonging to the clonal complex (CC)87 according to the Institut Pasteur nomenclature^19^. Using phylogenomic comparative analysis, Reid et al. point to a heightened role of ColV plasmid over the HPI in the evolution of this clone.

Thus, a clear picture of the respective role of the VAGs in the emergence of virulence, according to their genomic location and to the phylogenetic background of the strains, is lacking. In the present work, we deciphered the role of the ColV plasmid and HPI in the CC87 virulence using GWAS based on the mouse assay phenotype. We then extended the study of the role of the various iron capture systems to the species as a whole. Our goal was to understand how functional redundancy, epistatic interaction and genome location are acting at the species level.

## Results

### The CC87 is composed of five subgroups, one of them exhibiting numerous VAGs and antibiotic resistance genes

We first reconstructed the phylogenetic history of both ST58 and its sister group ST155 which are parts of the CC87 by analyzing the core SNPs of 232 strains, including 26 ST58 strains representative of the diversity previously described by Reid et al. (Figure 1, Table S1). These strains were recovered from humans (n=125), animals (n=87) as well as the environment (n=20) and were diverse in terms of geographical origin (Europe n=91, America n=59, Australia n=50, Africa n=30, and Asia n=2).

**Figure 1:**
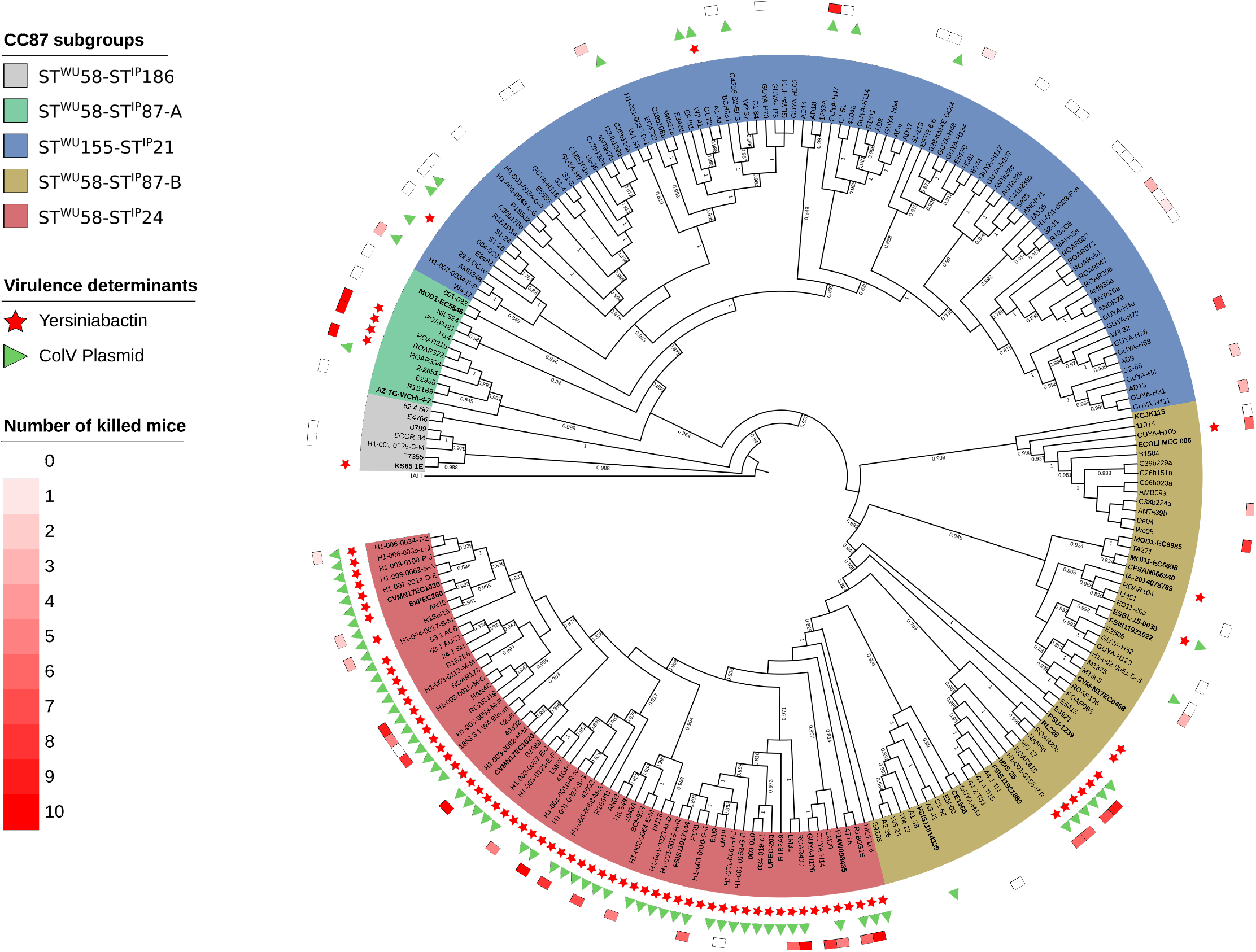
Maximum likehood core genome phylogenetic tree of the 232 B1 phylogroup CC87 (Institut Pasteur scheme numbering) strains. The five CC87 subgroups (ST^WU^58-ST^IP^186, ST^WU^58-ST^IP^87-A, ST^WU^155-ST^IP^21, ST^WU^58-ST^IP^87-B, ST^WU^58-ST^IP^24) based on the Warwick University and Institut Pasteur MLST schemes are highlighted in color. The presence of the high pathogenicity island (HPI) is highlighted by red stars and ColV Plasmid (defined as proposed by Reid *et al*.^18^) by green triangles. In the outermost circle, colored squares represent the number of mice killed over ten in the mouse model of sepsis^11^. The 26 ST58 genomes obtained from the study by Reid *et al*. are in bold. The tree is rooted on IAI1 (non-CC87, phylogroup B1). For the sake of readability, branch lengths are ignored and local support values higher than 0.7 are shown.

By rooting the tree on the B1 phylogroup ST1128 IAI1 strain, and based on both Warwick University (WU) and Institut Pasteur (IP) MLST schemes, five main subgroups were distinguished (Figure 1). The most basal one is the ST^WU^58-ST^IP^186 (ST58/186) subgroup (n=7), followed by the ST58^WU^-ST^IP^87-A (ST58/87A) (n=12), ST^WU^155-ST^IP^21 (ST155/21) (n=93) and ST^WU^58-ST^IP^87-B (ST58/87B) (n=57) subgroups, and the more recently emerged ST^WU^58-ST^IP^24 (ST58/24) (n=63) subgroup. Importantly, the latter encompasses the BAP2 cluster described by Reid et al.^18^

We then screened our collection for VAGs and antimicrobial resistance genes (ARGs). As reported previously^18^, the ST58/24 subgroup exhibited significantly more VAGs classified in toxin, protectin and iron acquisition system categories (Figure 2A, Table S2) and was predicted to be more antibiotic (ampicillin and trimethoprim) resistant (Figure S1, Table S3) than the other subgroups. Moreover, almost all strains in this subgroup carry the HPI and the ColV plasmids. Of note, whereas this subgroup represented 27% of our collection, it encompassed two-third of the strains isolated from human extra-intestinal infections (Figure S2), an over-representation also observed by Reid *et al*.^18^

Altogether, these first analyses confirmed that virulence and resistance genes within the *E. coli* B1 phylogroup ST58/ST155 clonal complex (CC87) are mainly present in the ST58/24 lineage.

**Figure 2:**
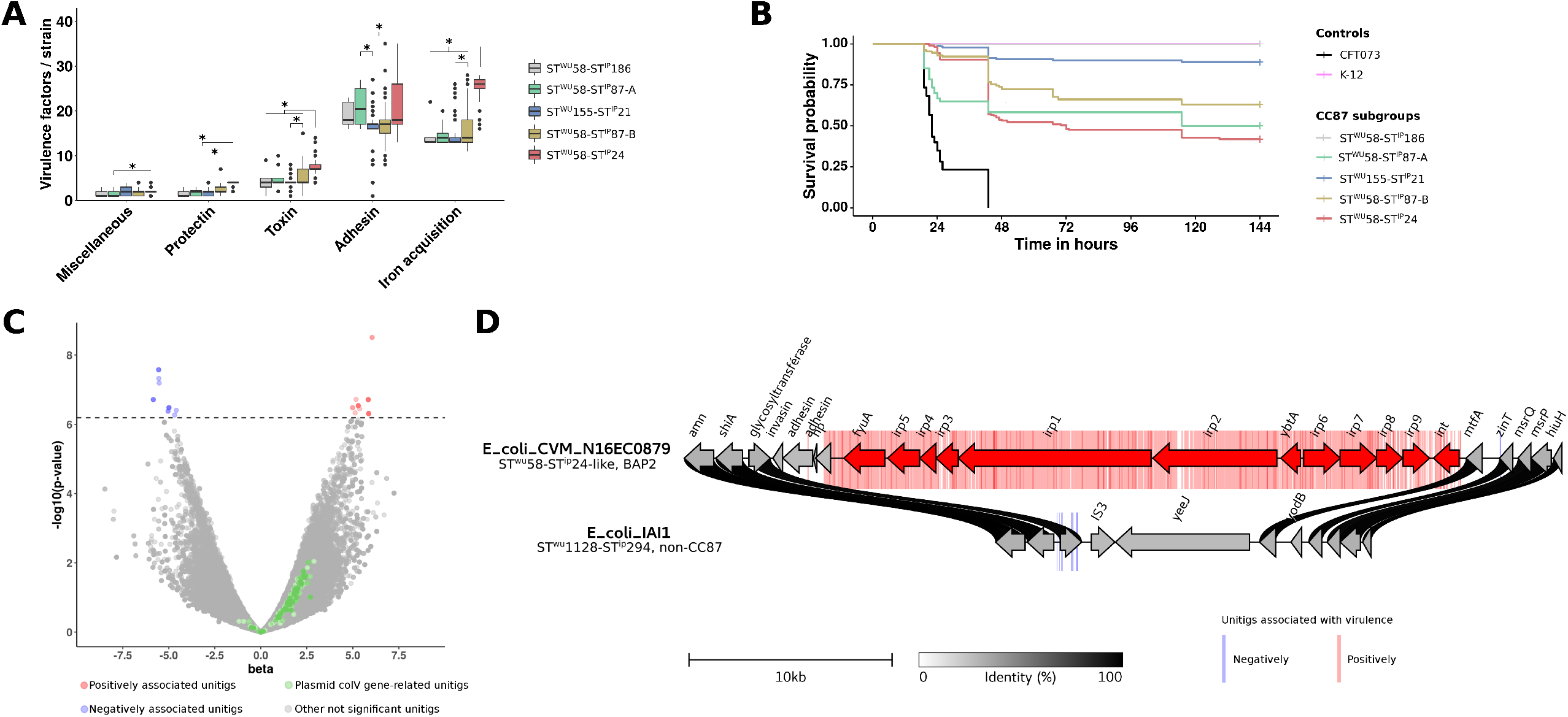
From genotypic and phenotypic characterization of CC87 extra-intestinal virulence to identification of genetic determinants by GWAS. A) Number of virulence-associated genes per strain among the six main functional classes of virulence according to CC87 subgroups. Significant differences are highlighted by asterisks. B) Kaplan-Meier survival curves of the mouse sepsis assay. Note that the pink (K-12) and grey (ST^WU^58-ST^IP^186) curves overlap. C) Results of the unitigs association with virulence. The p-value of the association is shown on the y-axis, the effect size (beta) on the x-axis and the significance level with a dotted line. The unitigs positively and negatively associated with the phenotype are highlighted in red and blue, respectively. The unitigs found in genes belonging to the ColV plasmid (as described by Reid *et al*.^18^) are highlighted in green. Other non-significant unitigs are in grey. D) Physical map of the genome region where significant association with the virulence in mice were observed. Unitigs positively or negatively associated are represented in the background of the map by red and blue bars, respectively. Genes positively associated with virulence are represented by red arrows and include the whole HPI. The fully sequenced and circularized genomes of *E. coli* CVM_N16EC0879 (ST^WU^58-ST^IP^24-like, BAP2 in the study by Reid *et al*.^18^) and IAI1 (ST^WU^1128-ST^IP^294, non-CC87) were used as reference. The links between the maps are colored according to amino acid identity.

### Mouse sepsis assay coupled to GWAS identifies the HPI as the major driver of virulence within the CC87

In a second step, we tested a set of 70 strains (30% of our data set) distributed in the five main subgroups in the mouse model of sepsis (Figure 1). Twenty-four strains have been tested previously^15^ whereas the 46 remaining were tested in the present work (Table S1).

Mouse survival curves showed that the strains exhibited different levels of virulence according to their subgroup belonging (Figure 2B). The ST58/24 subgroup killed more than all CC87 subgroups except the ST58/87A, in accordance with the VAG content. The ST58/186 and ST155/21 killed less than the other CC87 subgroups, ST58/186 even behaving as the K-12 avirulent control strain. Interestingly, the virulence observed for the B1 phylogroup strains is significantly lower than the control B2 phylogroup CFT073 strain, a potential effect of the role of the genetic background of the strains on the expression of virulence^13^.

To identify the genetic determinants responsible for the mouse virulence phenotype, we performed a GWAS with Pyseer^20^ using as input the unitigs calculated from the genome assemblies or the gene presence/absence and the number of mice killed by each strain. As shown on the volcano plot (Figure 2C), 417 and 16 unitigs were positively (in red) and negatively (in blue) significantly associated with the phenotype, respectively. All these unitigs except one correspond to the HPI insertion region (Figure 2D). The only non-HPI-related unitig was associated with a gene encoding a phage minor tail protein. Moreover, gene presence/absence also points to the HPI with 12 genes over the 13 significantly associated with virulence related to this virulence determinant (Figure 2D). In contrast, no unitig or gene related to the ColV plasmids were significantly associated with virulence (Figure 2C).

### At the species level, both *aer* and *sit* operons, but not the plasmid by itself, are involved in the mouse sepsis phenotype

We then wanted to evaluate the role of the VAGs used to infer the presence of ColV plasmids in extra-intestinal virulence, but this time at the species level. For this purpose, we used the data of a GWAS analysis performed on 370 strains of the genus *Escherichia* that have been tested in the mouse sepsis assay, including 24 strains of the CC87 (see above). In this work, the HPI was the strongest genetic element associated with virulence in mice, followed by the aerobactin and *sit* operons^15^.

We determined the genomic location (i.e. chromosomal or plasmidic) of the VAGs with PlaScope^21^. From this analysis, we were able to show that some genes were strictly plasmidic (*cvaC, cvi, etsABC, hlyF, ompT*) whereas others could be located on both elements. The VAGs *iucABCD* and *sitABCD* were predominantly chromosomal in 53.96-54.35 % and 72.73-73.27 % of the cases, respectively. The distribution of VAGs also differed according to the phylogeny (Figure 3). We found most of the VAGs in phylogroups B1, B2, C, D, F and G at a variable level. They were predicted to be almost exclusively located on plasmids in phylogroups B1, C and G, with a prevalence over 40% in the latter two phylogroups. At the opposite, chromosomal location was mainly observed in the archetypal ExPEC phylogroups B2, D and F. We ran the same analysis on the 232 genomes of the CC87 and confirmed the highest prevalence (70%) of VAGs in the ST58/24 subgroup and the almost exclusive plasmidic location of the VAGs in this population (Figure S3).

**Figure 3:**
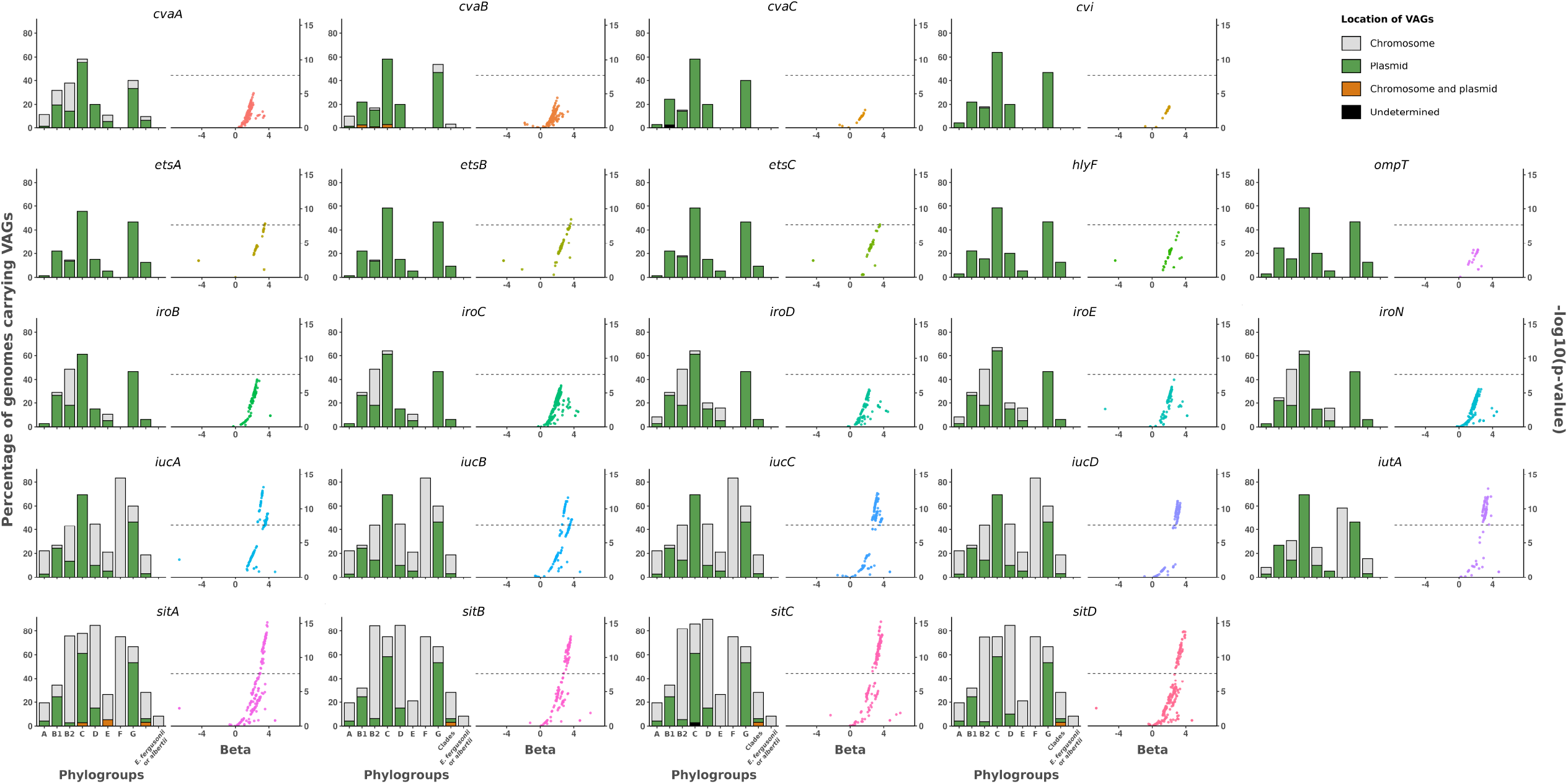
Prevalence and location of VAGs and association between virulence and unitigs within these VAGs among 370 genomes of the *Escherichia genus*. For each VAG two plots are represented. The bar plot represents the prevalence according to the phylogroup/genus and the predicted location. Plasmidic location is highlighted in green, chromosomal location in grey and cases with both chromosomal and plasmidic location in orange. Location of VAGs could not be determined in two genomes (*cvaC* in phylogroup B1 and *sitC* in phylogroup C) and is shown in black. The whole dataset is composed of genomes of strains from phylogroup A (n=72), B1 (n=41), B2 (n=111), C (n=36), D (n=20), E (n=19), F (n=12), G (n=15), clades (n=32) and *E. fergusonii* and *E. albertii* (n=12). The scatter plot represents the association between unitigs within the VAGs and the virulence in mice. The p-value is shown on the y-axis, the effect size (beta) on the x-axis and the significance level of the GWAS analysis with a dotted line.

Then, using GWAS, we looked at associations of unitigs belonging to each of these VAGs with virulence in mice. Interestingly, we found significant associations mainly for the aerobactin and *sit* operons (Figure 3). The sequences of VAGs do not significantly differ depending on their predicted location (Figure S4) and when analyzing the association results according to the predicted location of each VAG in each strain, we did not observe appreciable differences according to genome localization (Figure S5). When focusing on genes that were exclusively located on a plasmid, such as *hlyF* and *ompT*, we found that all unitigs mapped to them were far below the significant threshold (Figure S5). If we consider the presence of the ColV plasmid as a covariate in the GWAS analysis, only the unitigs associated with *iucABCD/iutA* and *sitABCD* remain significant, arguing for a role beyond their simple localization on the plasmid (Figure S6).

Overall, these data indicate that within the *Escherichia* genus, the sole presence of the ColV plasmids has little or no role in virulence in the mouse model, with the exception of the two aerobactin and *sit* operons. However, the predominant location of the latter is in fact chromosomal, reinforcing the idea that ColV plasmids are not themself the main driver of virulence.

### The HPI needs other VAGs to express full virulence

Next, we assessed the respective role of the iron acquisition-related operons *aer, sit* and *iro* as compared to the higher predictor of mouse death, the HPI, in the same 370 strains dataset. The presence of the HPI was associated with a significantly higher number of mice killed per strain in all cases except strains with *iro* alone, *iuc* alone and the combination *iuc/iro* (Figure 4). However, the presence of the HPI alone without *iro, aer* and *sit* operons in a strain was associated with the death of mice (at least 2 mice killed over 10) in only 50% of the cases. The avirulent strains despite the presence of the HPI belonged to phylogroups A (n=5/9), B2 (n=2/9), C (n=2/9) (Figure S7). Moreover, when we compared strains carrying the HPI depending on the presence of the other operons, the presence of the HPI alone was associated with significantly fewer mice killed per strain than in the cases of the HPI with *sit, sit/iro, sit/iuc* or *sit/iuc/iro*. No differences were observed for strains carrying only *iro, iuc* or *iro/iuc* probably due to the limited number of strains.

**Figure 4:**
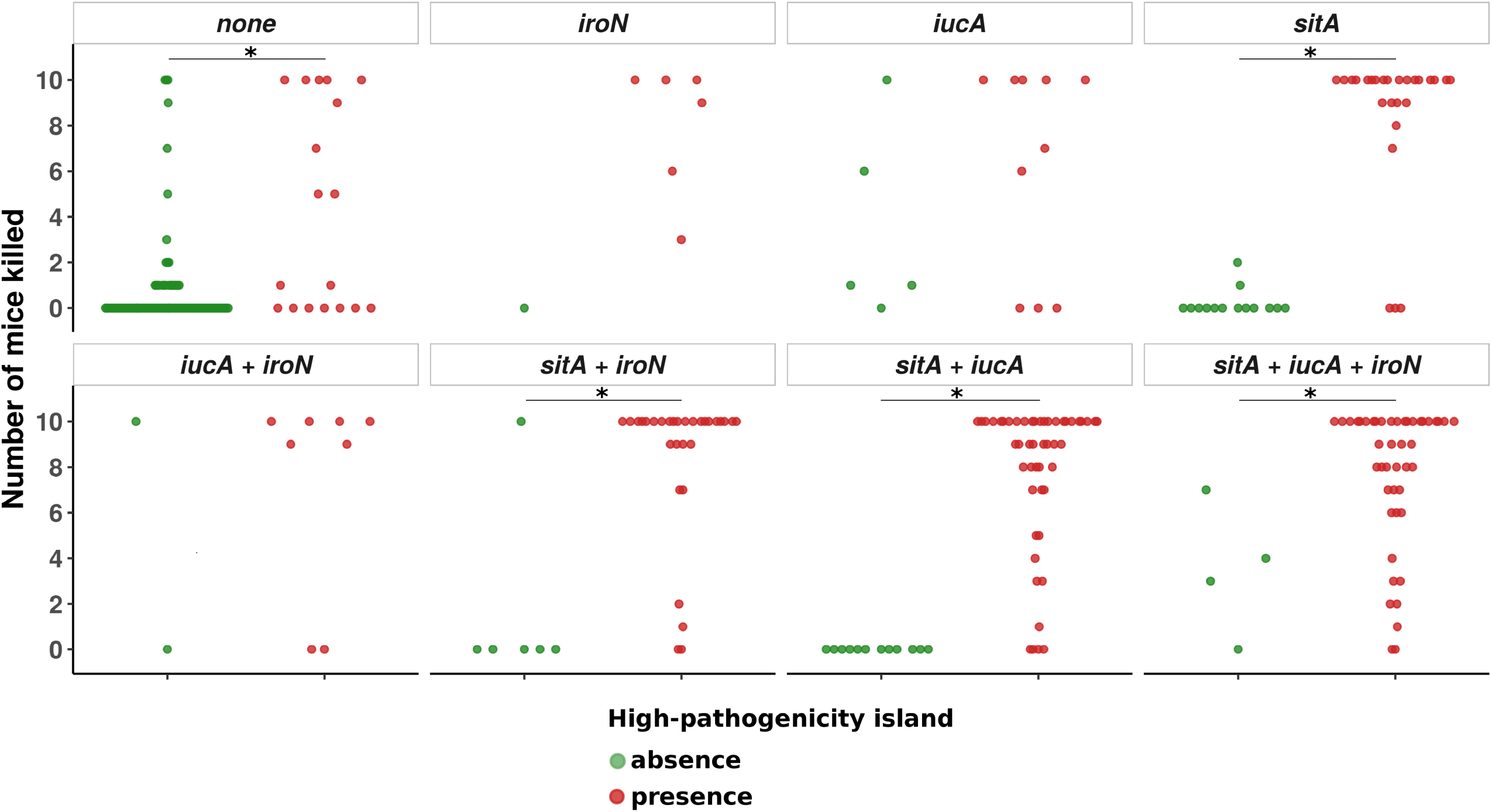
Number of mice killed over ten according to the combination of iron acquisition-related VAGs *iroN, iucA, sitA*. In each facet, each point represents the number of mice killed by a given strain according to the VAG or combination of VAGs it carries. Strains with or without the HPI are represented by red and green points, respectively. Significant differences are highlighted by asterisks. A related figure detailing the phylogroup of each strain is available in Figure S7.

Overall, these data indicate that the presence of HPI is necessary but not always sufficient for a strain to be virulent, with other iron uptake systems required to express full virulence. Conversely, these accessory systems alone are not sufficient to acquire full virulence.

### The HPI co-occurs with *aer, sit* and *iro* operons in *E. coli* genomes at different frequencies depending on the phylogeny of the strain

To get a broader view of VAG co-occurrences and their genomic localization, we used the 2,302 high-quality circularized *E. coli* genomes available in RefSeq (https://www.ncbi.nlm.nih.gov/refseq/). Using this dataset prevented us from the potential bias resulting from the Illumina short-read sequencing when we analysed the genomic location of VAGs among the 370 *Escherichia* strains. Nonetheless, we confirmed that PlaScope enables a classification of chromosomal and plasmidic contigs with high performances (Figure S8).

First, we focused on the five main ST responsible for bacteremia in France^22^ as well as the STc58 to quantify co-occurrence frequencies and prevalence of VAG pairs. In all these ST/STc, except the ST73, the plasmidic genes co-occurred at a similar prevalence (from few percent in ST10 and ST131 to 75% in ST95) which supports the frequent co-location of these VAGs (Figure 5). Moreover, these plasmidic VAGs usually occurred in strains which carry *fyuA*, except in the commensalism-associated ST10. More globally, we can see specific patterns. In the case of ST131, the overall pattern is dominated by a very high prevalence and frequent co-occurrence of chromosomal VAGs, with a notable absence of *iroN*. This chromosomal dominant pattern is exacerbated in the ST73 with an almost complete fixation of VAGs on the chromosome. The ST69 and ST95 both present high prevalence and co-occurrence frequencies of chromosomal and plasmidic VAGs, without chromosomal *iroN* or *iucA*, respectively. Of note, this is in agreement with the ColV plasmids frequently observed in the ST95^23^. On the opposite and consistent with its commensal behaviour, the ST10 carries few VAGs, which are usually clustered on plasmids. Finally, for the STc58, in accordance with our previous observation and data from Reid *et al*.^18^, *iroN, iucA* and *sitA* co-occurred mainly and frequently on plasmids and are usually associated with the HPI.

**Figure 5:**
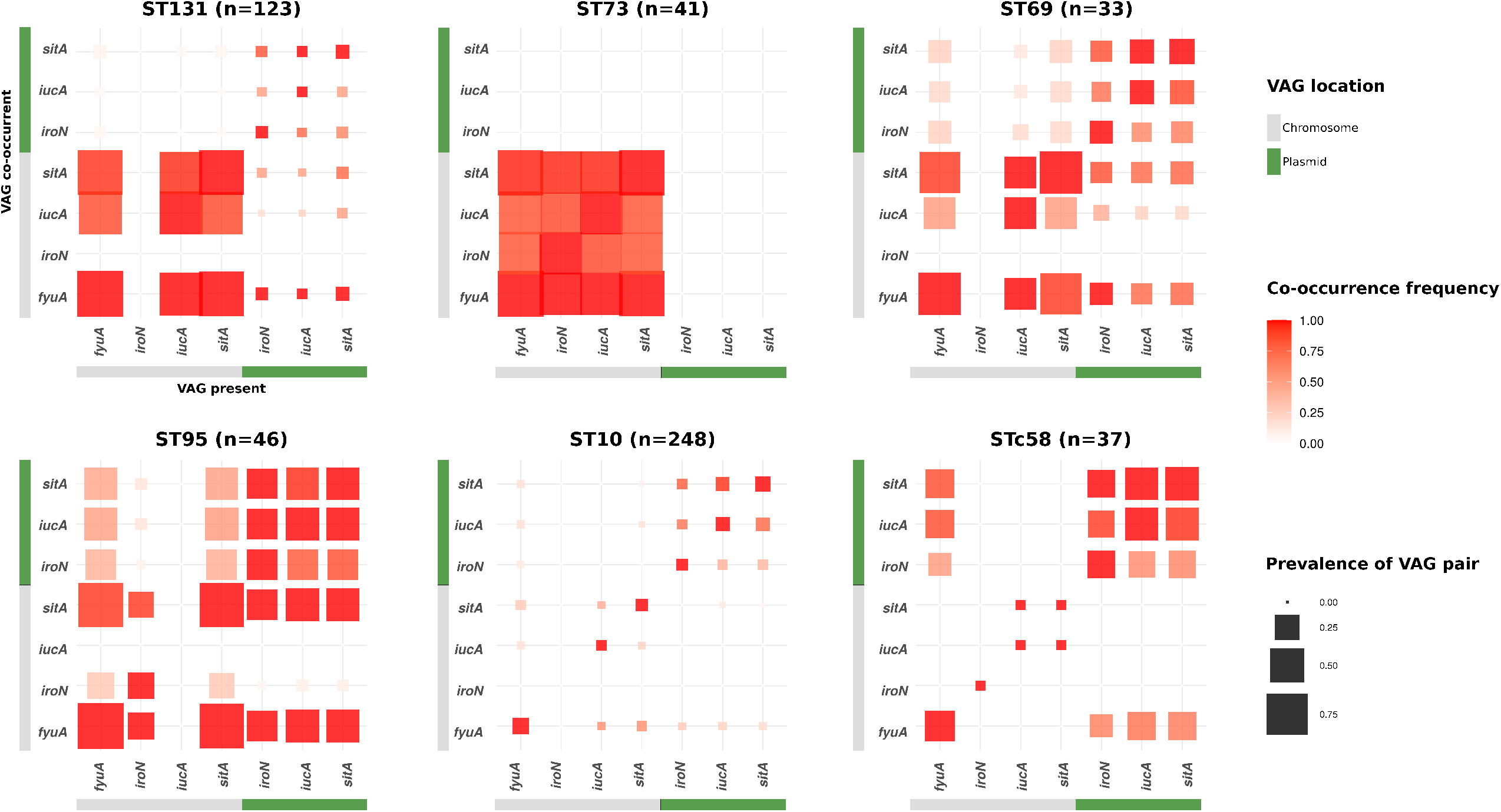
Co-occurrence frequency and prevalence of iron acquisition-related VAG pairs among 528 fully circularized *E. coli* genomes belonging to ST131, 73, 69, 95, 10 and STc58. For each given VAG on the x-axis, the frequency of co-occurrence with the VAGs on the y-axis is highlighted by a colour gradient. The size of each square is proportional to the prevalence of the VAG pair in the given ST/STc. VAGs are separated according to their location on the chromosome in grey or on the plasmid in green. The co-occurrence frequencies of the VAGs and the prevalence at the phylogroup scale are available in Figure S9.

Then, looking at the broader scale of phylogroups, the patterns of co-occurrence are also phylogroup specific with mainly three kinds of pattern (Figure S9). In phylogroups A, B1 and E, very few co-occurrences were found, with the exception of plasmidic VAGs, although their prevalence was low. In phylogroups C, F and G, we observed mainly frequent co-occurrences and high prevalences of plasmidic VAGs. The latter frequently co-occurred with *fyuA*, ranging from 41.7% to 81.3%. Finally, among phylogroups B2 and D we commonly found co-occurrences of chromosomal VAGs that are also highly prevalent. Moreover there were also strong co-occurrences between plasmidic VAGs and chromosomal *sit* and HPI. Of note, plasmidic VAGs frequently co-occurred with the HPI in phylogroups B2, C, D, F and G. But the reverse was true only in groups C, F and G.

Overall, the co-occurrences of VAGs show patterns that are specific to ST/STc and/or phylogroups. Frequent co-occurrences and high prevalences of chromosomal VAGs are mainly observed in typical ExPEC like ST131, 73, 95, 69 or more broadly in phylogroups B2 and D. Of note, *iro* or *iuc* may be absent from the chromosome depending on the clone. Interestingly, plasmidic *iro, iuc* and *sit* operons mainly arise in strains that already carry the HPI. But the reverse is usually not true with the exception of phylogroups C, F and G.

### The VAG co-occurrences result from selection

To determine whether these co-occurrences were the result of chance and timing or selection, we search for the sites of chromosomal insertions of the genes. From this analysis, we could expect either a single site if the arrival is due to chance or multiple sites indicating evolutionary convergence through multiple arrivals. Whereas the HPI was mostly inserted at a single site but spread by homologous recombination within the *E. coli* species^17^, *iro, sit* and *iuc* genes were inserted in multiple chromosomal sites (6, 14 and 10, respectively) (Table S4, S5, S6). These sites were mainly ST specific but unrelated to the strain phylogeny, the same site being found in different phylogroups. Very rarely, several sites were observed within a single ST as in ST131.

Then, to get additional evidence for selection, we compared the phylogenetic histories of *aer, sit* and *iro* based on nucleic sequence alignments of operons for each location (i.e. chromosome or plasmid) to the strain phylogeny, incongruences between gene and strain phylogenies indicating multiple horizontal gene transfer events (Figure S10). When comparing the phylogeny at the phylogroup level, a very high level of incongruence was noted for the plasmidic genes (Figure S10 B, D, F) and to a lesser extent for the chromosomal genes (Figure S10 A, C, E). We then compared patristic distances between all possible pairs of genomes according to their location and ST and/or phylogroup (Figure 6). From this analysis, we noted as expected that the patristic distances between plasmid operons were in all cases very small, reflecting the very high mobility of plasmid sequences with homogenization (Figure 6B, D, F). But we observed also that despite a global increase of patristic distances in different ST and/or phylogroup, some operon sequences from strains of different STs or phylogroups show patristic distances as small as between strains of the same ST (Figure 6A, C, E), highlighting the existence of some mobility of the chromosomal sequences.

**Figure 6:**
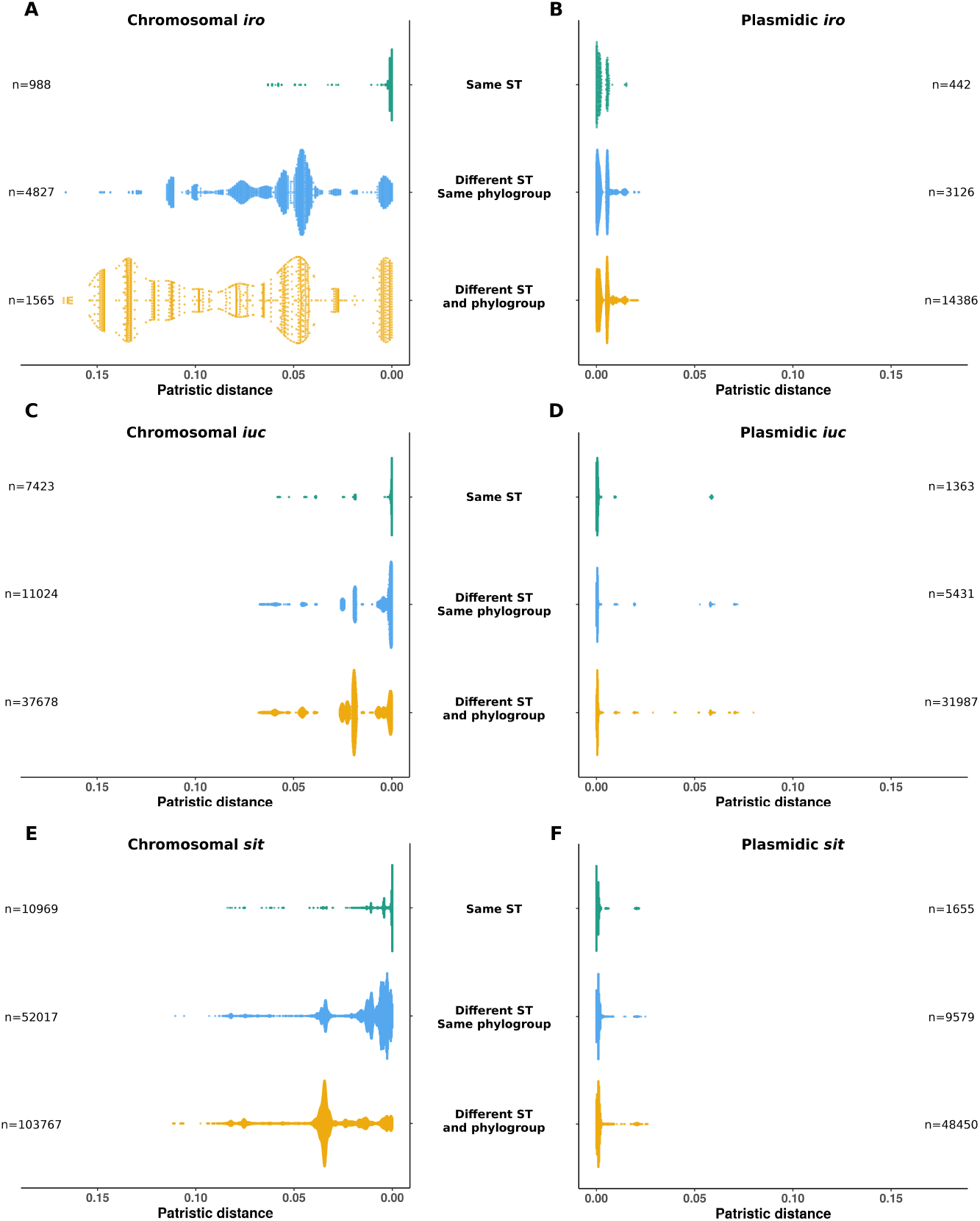
Distributions of patristic distances between *iro, iuc* and *sit* operons according to their location and to the sequence types and phylogroups of the strains. The scatter plots represent the patristic distances between pairs of (A) chromosomal *iro* operons, (B) plasmidic *iro* operons, (C) chromosomal *iuc* operons, (D) plasmidic *iuc* operons, (E) chromosomal *sit* operons and (F) plasmidic *sit* operons. The number of pairs of genomes involved in each category is specified opposite each scatter plot. For the sake of readability, the highly divergent sequences of chromosomal *iuc* from genomes GCF_010725305.1 and GCF_002946715.1 and chromosomal *sit* from genome GCF_024225755.1 were not included.

In sum, we found multiple arrivals of VAGs carried by plasmids and to a lesser extent at the chromosomal sites within the species phylogenetic history, a hallmark of selection.

## Discussion

Classically, the confirmation of the role of a VAG in virulence is obtained by inactivating the gene and by showing that the KO mutant is not anymore virulent or is outcompeted by the wild type strain in co-infection assays. Ideally, the complementation of the mutant should restore the wild-type phenotype. Such an approach has proven to be very powerful but it is particularly time consuming and usually only allows for the testing of single or double mutants^24–29^. Mutants of the seven pathogenicity islands (PAIs) of the B2 phylogroup *E. coli* 536 strain alone or in combination have been constructed, removing up to 7.7% of the genome and allowing to test simultaneously blocks of hundreds of genes^30,31^. The arrival of rapid and inexpensive genome sequencing has shaken up these classical approaches by enabling large-scale epidemiologic studies on bacterial virulence phenotypes^32^. These studies infer evolutionary scenarios leading to functional consequences, such as emergence of virulence, based solely on comparative phylogenomic data^18,33–37^. Such sequence data can also be used in GWAS if they are linked to virulence phenotypes^15,38^.

The picture emerging from these studies is that extra-intestinal virulence in *E. coli* results from a multigenic process depending of the phylogenomic background, with a major role of the iron capture systems^6^. However, little is known about the relative contributions of the VAGs according to specific lineages.

In the present work, we explored further the emergence of a specific extra-intestinal virulent lineage within *E. coli* ST58/CC87^18,19^. Interestingly, it belongs to the B1 phylogroup which usually encompasses mostly commensal and intra-intestinal pathogenic strains^6^. We showed, using a mouse model of sepsis together with a genetic association analysis, that the virulence phenotype among CC87 is strongly correlated to the presence of the HPI, regardless of the presence of the ColV plasmid. This is in disagreement with the work of Reid et al, who have proposed a somewhat direct role of this plasmid in *E. coli* virulence based only on in silico data^18^. In addition, we did not evidence at the *Escherichia* genus level any role of the ColV plasmid except by the fact that it is bearing the *aer* and *sit* operons. The latter are also frequently and even predominantly found on the chromosome as in the ExPEC B2, D and F archetypes which are highly virulent in mice^15^. This plasmid’s association in the history of ExPEC, as tested in the mouse sepsis assay, might therefore be restricted to carrying these genes in some sequence types. However, it can be hypothesized that the other genes could be involved in other aspects of extra-intestinal virulence or in commensalism.

As iron capture systems are key elements in the mouse sepsis assay, we wanted to understand their respective role and the evolutionary forces acting on them. In addition to the HPI, *sit* and *aer* operons, we took in consideration *iro* operon that is also present on the Col plasmids and on the chromosome. Our data show a hierarchy among the VAGs with the HPI having the preeminent role but the other operons potentializing the HPI effect (Figure 4). This is possibly true for *iro* genes for which we did not evidence a role by the GWAS (Figure 3). This hierarchy could be the result of complementarity between different functions encoded by the different operons, aerobactin, salmochelin and yersiniabactin being hydroxamate, catecholate and mixed-type siderophore, respectively^26^ whereas Sit is an iron and manganese transporter^29^. Interestingly, functional redundancy and preeminent role of specific iron acquisition systems have been observed for ExPEC in a urinary tract infection mouse model, allowing niche specificity within the urinary tract^26^. It has also been observed that salmochelin and aerobactin contribute more to virulence than heme in a chicken infection model^39^ and have a cumulative effect^9,39^. Discrepancies between these studies and our own data could result from the fact that (i) the HPI was not considered, (ii) the results were based on single strain KOs and (iii) animal models other than mice were used. Thus, both hierarchy and cumulative effect of iron capture VAGs seem a common theme in extra-intestinal virulence. In addition, the importance of the tested systems has been shown to depend on the strain used^14,40^. This could be due to different associations of systems present in the strains.

We provide strong evidence that the acquisition of these VAGs is under strong selection as they arrived multiple times via a plasmid and to a lesser extent within the chromosome. A striking feature of our work is the ST (phylogroup) specific patterns of co-occurrences (Figure 5). The Col plasmid is present in ST95, 69 and 58, rare in the ST131and totally absent in the ST73. Mirroring this pattern, the chromosomal VAGs are highly prevalent in ST73 and 131 but in the latter without *iro*, and present in ST95 but without *aer*. Similar lineage-specific co-occurrence patterns of iron related VAGs have been recently shown in *Klebsiella* spp. isolates^41^ (nat mic). Interestingly, these patterns are niche specific (human clinical versus porcine isolates) and resulted from multiple arrival events. Such a marked pattern (*i.e*. on/off) indicates that very strong antagonism at the genome level (plasmid vs chromosome) and/or between the end products of the genes exists. A potential explanation of this antagonism is the presence of epistatic interactions at the genome level. The role of the strain genetic background in the acquisition/maintenance of acquired genes has been recently documented for the CRISPR-Cas systems acquired by horizontal gene transfer^42^ by showing inhibition of non-homologous end-joining repair system by Csn2 Cas protein from which encoding genes never co-occurred^43^.

Our work has some limitations. First, our CC87 data can suffer from a lack of power due to the small number of CC87 mouse tested strains. Our analysis does not exclude minor roles of other loci such as the other ColV plasmid genes. However, due to ethical considerations, it is difficult to increase the number of mice used as we already obtained a strong signal. Second, although a well-accepted model, our mouse sepsis assay cannot mimic the invasion step of the infection as we inoculate the bacteria directly in the neck, which is a non pathophysiogical process.

Nevertheless, we bring strong evidence that a cumulative effect of iron capture systems is involved in *E. coli* extra-intestinal virulence with the HPI being the major player. The prevalence, the co-occurrences and the genomic location of these VAGs depend of the phylogenetic lineage, supporting the role of epistasis in the emergence of virulence. Deciphering the molecular mechanisms involved will be a real challenge for the future years.

## Methods

### *E. coli* CC87 dataset

The collection is composed of strains belonging to the CC87 (Institut Pasteur scheme numbering) which gathered strains from both ST58 (n=139) and its sister group ST155 (n=93), all from phylogroup B1. The strains were sampled from humans (n=125), domestic (n=66) and wild (n=21) animals and environment (n=20) (Table S1). Human strains corresponded to commensals (n=64), ExPEC (n=60) and InPEC (n=1) whereas animal strains were all commensals (Figure S1). The geographic origin was diverse (Europe n=91, America n=59, Australia n=50, Africa n=30, and Asia n=2). Of note, 26 ST58 strains previously described by Reid *et al*.^18^ were also included. All our strains were sequenced on Illumina platforms. Genomes were annotated with Prokka using standard parameters^44^. Bioprojects and accession numbers of the genome are detailed in Table S1. Fasta and gff files are available on figshare^45,46^.

### Genome typing, VAGs/ARGs screening and annotation

We determined multi-locus sequence types (MLST) of each CC87 strain using mlst ^47,48^ based on both the Warwick University^49^ and the Pasteur Institute schemes^50^. The O-type, H-type and fimH of the strains were determined with Abricate^51^ with 90% identity and 90% coverage based on ecoh^52^, serotypefinder^53^ and fimTyper^54^ databases. We searched for virulence associated genes (VAGs) and antibiotic resistance genes (ARGs) as previously described using Abricate with 90% identity and 90% coverage^22^. Our virulence database was composed of VirulenceFinder^55^, VFDB^56^ and specific genes from extra-intestinal *E. coli*. The VAGs were classified into 6 main families, namely invasin, protectin, toxin, adhesin, iron acquisition, and miscellaneous. We also searched for point mutations responsible for betalactam and fluoroquinolone resistance using pointFinder^57^. Based on ARGs and mutation presence/absence we predicted resistance phenotype to various antibiotics: ampicillin, piperacillin-tazobactam, cefotaxim/ceftriaxone, cefepim, carbapenems, fluoroquinolones, gentamicin, amikacin, sulfamides, trimethoprim, chloramphenicol, tetracyclines, colistin (Table S3).

### ColV plasmid inference and plasmidic sequences prediction

To infer the presence of ColV plasmids we searched for VAGs from 6 gene sets (*cvaABC/cvi, iroBCDEN, icuABCD/iutA, etsABC, ompT/hlyF, sitABCD*) using Abricate with 90% identity and 95% coverage as proposed by Reid et al.^18^. Gene sequences were retrieved from the plasmid pAPEC-O2-ColV (RefSeq: NC_007675.1). Then, from these results, we inferred the presence of ColV plasmids on the basis of Liu’s criteria^58^, i.e. if at least one gene from 4 of the 6 gene sets was present. We also predicted chromosomal and plasmidic sequences using PlaScope^21^, which is a targeted approach to classify contigs based on a database of chromosome and plasmid sequences of *E. coli* (database available on Zenodo^59^).

### Core-SNP phylogeny of the CC87

We built a core-SNP phylogeny by aligning the CC87 genomes to the reference strain IAI1 (ST^WU^1128-ST^IP^294, non-CC87, phylogroup B1) using Snippy 4.4.0^60^ and we filtered recombination with gubbins v2.3.4^61^ using standard parameters. The recombination free alignment was used to compute a phylogenetic tree with FastTree v2.1.11^62^ with the generalised time reversible (GTR) + gamma substitution model. Phylogenetic tree was annotated with Itol^63^.

#### Mouse model of sepsis

We assessed the intrinsic virulence of 70 CC87 strains using a mouse model as previously described^11^. In this model, 10 female mice OF1 of 14–16 g (4 week-old) from Charles River (L’Arbresle, France) are inoculated with 10^8^ *E. coli* cells subcutaneously in the neck and monitored for six days. Two control strains (CFT073 and K-12) killing 10 and zero mice over 10, respectively, were systematically added in each experimental series. The number of mice killed per strain is available in Table S1. Of note, 24 strains have been tested in a previous work^15^ in the same way, while the other 46 were tested in the present study (Table S1). In total this corresponds to a representative sample of each CC87 subgroup: ST^WU^58-ST^IP^186 (n=2/7), ST^WU^58-ST^IP^87-A (6/12), ST^WU^155-ST^IP^21 (28/93), ST^WU^58-ST^IP^87-B (13/57), ST^WU^58-ST^IP^24 (21/63). The protocol (APAFIS#4948) was approved by the French Ministry of Research and by the Ethical Committee for Animal Experiments, CEEA-121, Comité d’éthique Paris-Nord.

#### Genome-wide association study to identify virulence determinants

We performed a GWAS based on the results of the mouse model of sepsis to identify genetic determinants associated with virulence in the CC87. We ran the analysis with Pyseer using the unitigs calculated from the CC87 genome assemblies (unitig-caller 1.2.1) or the gene presence/absence calculated with Roary 3.12.0^64^ and the number of mice killed by each strain (continuous phenotype from 0 to 10). Unitigs are a compact representation of the pangenome in the form of extended DNA sequences of variable length^65^. The population structure was taken into account using the FastLMM linear mixed-model^66^ and the patristic distances between each pair of strains extracted from the recombination-free phylogenetic tree previously computed. The p-value threshold for each analysis (i.e. unitigs and gene presence/absence) was corrected using Bonferroni method considering the number of unique variant patterns as the number of multiple tests. These corrections resulted in p-values of 6.09E-07 and 1.56E-05 for unitigs and genes, respectively. Finally, to identify their location, we mapped back the statistically significant unitigs using bwa 0.7.17^67^ and bedtools 2.30.0^68^ to 3 fully sequenced reference genomes of *E. coli:* CVM_N16EC0879 (ST^WU^58-ST^IP^24-like, phylogroup B1, GenBank: CP043744.1), IAI1 (ST^WU^1128-ST^IP^294, non-CC87 phylogroup B1, Refseq: NC_011741.1), S88 (ST^WU^95-ST^IP^1, phylogroup B2, Genbank: CU928161.2). The coordinates of the significant unitigs and genes were used to draw physical maps of the region of interest from the reference genomes CVM_N16EC0879 and IAI1 using Clinker^69^ and geom_segment from ggplot2^70^. Input files and raw results of the GWAS analysis are available on figshare^71–75^.

#### Role of VAGs from ColV plasmid in virulence at the species level

To assess the role of each VAG used to infer the presence of colV plasmids, we retrieved the genomes of 370 strains representative of the diversity of the *Escherichia* genus as well as the results of the GWAS previously done on these strains^15^. First, we screened the genomes for ColV plasmid VAGs as described above and classify contigs of each genome using PlaScope. Then, we constructed a blastN database from nucleic sequences of all ColV plasmid VAGs found among the 370 genomes. We ran a blastN alignment using this database and the unitigs as query and considered only perfect matches (i.e. 100% identity over 100% of the unitig length). From this analysis we were able to associate each unitig and its p-value/beta-value to a given VAG. We also performed a blastN alignment of all sequences of a given VAG depending on its predicted location to compare VAGs sequences within and between different locations (i.e. plasmids and chromosomes) (Figures S4, S5).

#### Distribution and location of VAGs at the species level among fully sequenced genomes of *E. coli*

To analyse more thoroughly the distribution of VAGs used to infer ColV plasmids, we determined their prevalence among the 2,302 genomes of *E. coli* (taxid 562) available on Refseq on September 19^th^ 2022. Both fasta and genbank files were retrieved using ncbi-genome-download^76^. Accession numbers of assemblies and replicons (chromosomes and plasmids) are available in Table S7. ColV plasmid VAGs were screened as described above using Abricate. Then, for each couple of VAG/location in a given genome, we searched for other co-occurrent VAGs. Results are presented as heatmaps and colored according to the frequency of the co-occurrences and prevalence of VAG pairs.

In a second step, we determined the site of insertion in the cases of chromosomal insertion of VAGs. To this end, we performed a pangenome analysis of the 2,302 genomes, using Ppanggolin v1.2.74^77^ and the annotations from RefSeq as input. The gene families were computed by clustering protein sequences with 90% identity and 90% coverage. Then, we built the pangenome graph and partitioned the pangenomes. Finally, we used PanRGP^78^ to determine the regions of genome plasticity (RGP), which can be considered as genomic islands in the cases of chromosomal insertions. The RGP containing *iro, iuc* and *sit* operons were identified using the function “align”. We extracted the sequences and annotations of these RGP as well as 3000 pb upstream and downstream to determine the sites of chromosomal insertion. The pangenome file from Ppanggolin/PanRGP is available on figshare^79^.

Third, we also took advantage of this high-quality dataset to check for the performances of PlaScope. We analyzed the 2,302 fully sequenced genomes with this approach and compared the replicon assignments to those of RefSeq (Fig S8).

#### Phylogenic incongruence of VAGs among the 2,302 genomes of RefSeq

We extracted the nucleic sequences of three operons involved in iron acquisition: *iroBCDEN, iucABCD* and *sitABCD*. We aligned the sequences of these operons according to their location (i.e. chromosomal or plasmidic) using Mafft v7.310^80^ and computed phylogenetic trees with FastTree v2.1.11^62^ with the GTR + gamma substitution model. From these trees, we computed patristic distances between all possible pair of sequences according to their location and ST/phylogroup using “cophenetic” from the R package ape^81^. The trees were also annotated and visualized with Itols^63^.

#### Statistical analysis

The number of VAGs per strain was compared between the CC87 subgroups using Kruskal-Wallis test. Mouse survival curves were compared with the log-rank test between the CC87 subgroups and the control strains. The number of predicted resistant strains among each CC87 subgroup was compared with Fisher exact test. The number of mice killed per strain according to the presence of the HPI and to the presence of VAGs or combination of VAGs was compared by a one-way ANalysis Of Variance (ANOVA). When multiple tests were performed, the p-values were corrected using the Bonferroni correction method. All tests were two-sided with a 5% type I error. The results of statistical analysis are available in Table S2.

## Acknowledgements

We are grateful to Laurence Armand and Milen Milenkov for the access to the Madagascar strains and to Lucile Vigué for fruitful discussions on epistasis.

This work was partially supported by the “Fondation pour la Recherche Médicale” Equipe FRM 2016, grant number DEQ20161136698 to ED. MG was funded by the Deutsche

Forschungsgemeinschaft (DFG, German Research Foundation) under Germany’s Excellence Strategy - EXC 2155 - project number 390874280.

## Competing interests

The authors declare they do not have any competing financial and/or non-financial interests in relation to the work described.

## Author contributions

GR, OC and ED concepted and realized the research. BC helped in the manipulations of the sequences. MG was involved in the analysis of the GWAS data and the editing of the paper. GR and ED wrote the paper.

## Notes

### Competing Interest Statement

The authors have declared no competing interest.

