## Supplementary online material for "High pathogenicity island, *aer* and *sit* operons: a “ménage à trois” in *Escherichia coli* extra-intestinal virulence"

**Supplementary tables**

**Table S1**: Principal characteristics of the 232 Escherichia coli CC87 strains studied.

**Table S2**: Kruskal-Wallis test p-values after Bonferroni correction of VAGs, Log-rank test p-values after Bonferroni correction of the mouse survival curves, Fisher exact test p-values after Bonferroni correction of antibiotic resistance data, ANOVA test p-values for the comparison of number of mice killed depending on VAG combinations.

**Table S3**. Antibiotic resistance prediction according to genes/mutations.

**Table S4**. Chromosomal insertion sites of *iroN*.

**Table S5**. Chromosomal insertion sites of *iucA.*

**Table S6.** Chromosomal insertion sites of *sitA.*

**Table S7.** List of the 2302 fully sequenced genomes from RefSeq.

**Supplementary Figures**

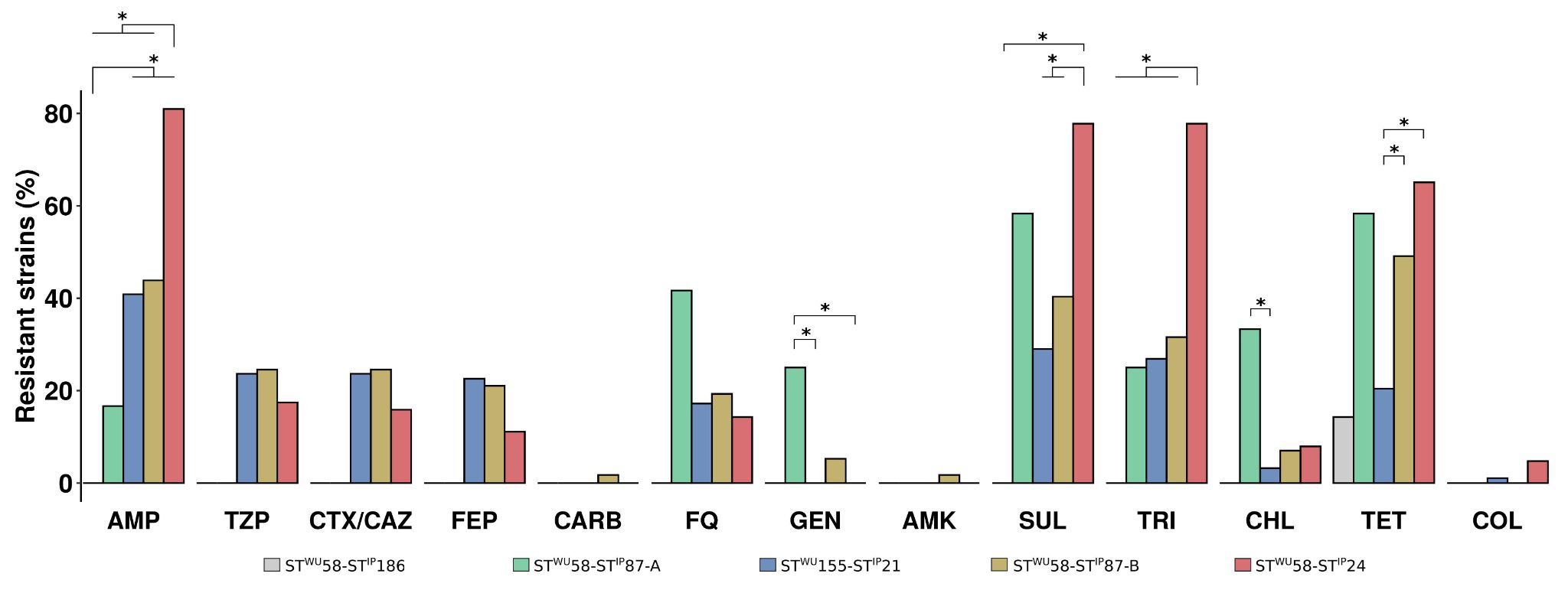

**Figure S1:** Predicted resistance phenotypes of the strains. The results are presented as the percentage of resistant strains for thirteen antibiotics of clinical and/or veterinary importance. Bars are coloured according to the CC87 subgroups. We predicted the phenotypes from the presence of genes or mutations in genomes as in Royer *et al.*^1^ and based on the genotype to phenotype predictions described in **Table S3**. Significant differences are highlighted by asterisks.

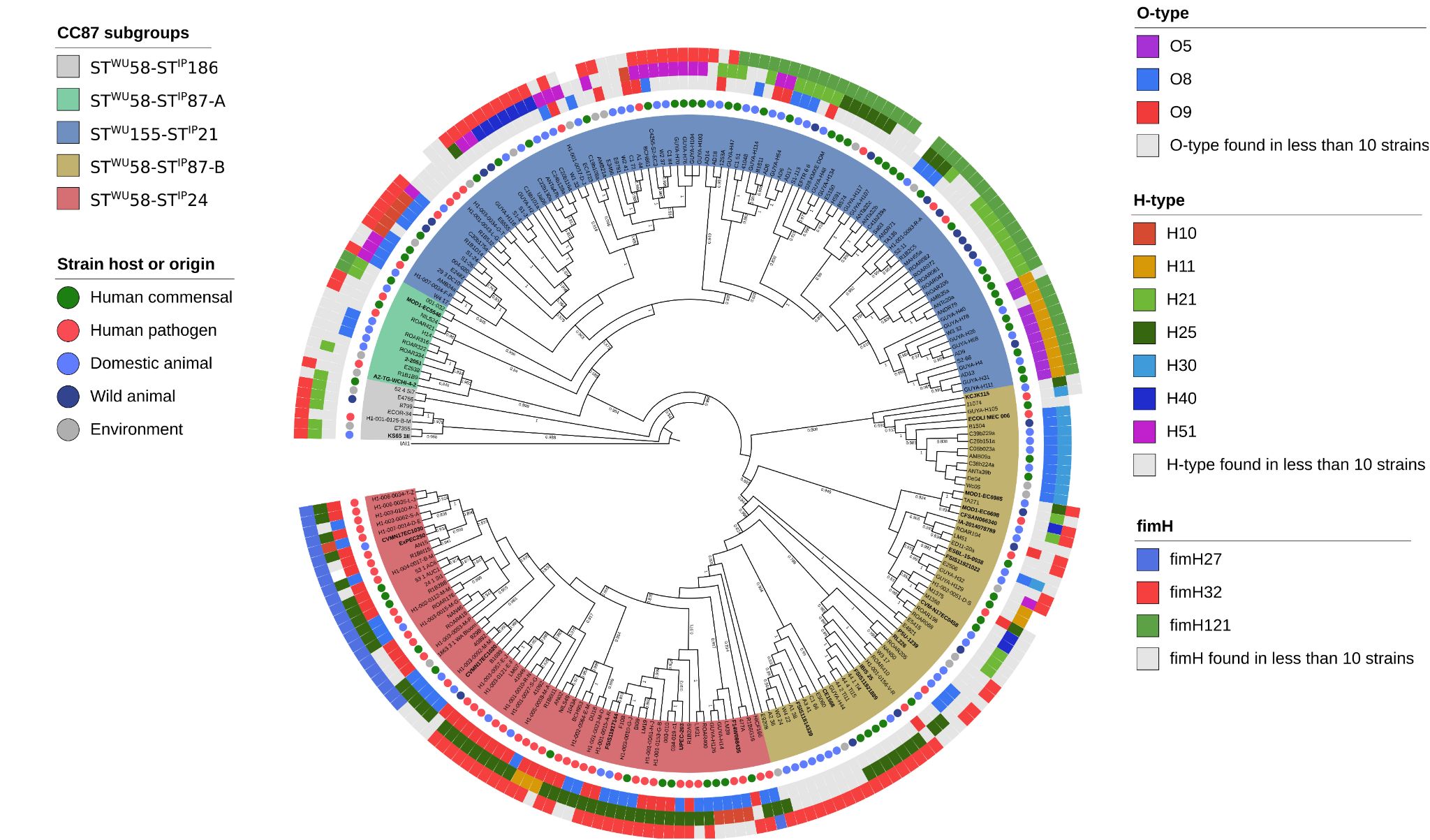

**Figure S2**: Maximum likelihood core genome phylogenetic tree of the 232 B1 phylogroup CC87 (Institut Pasteur scheme numbering)^2^ strains. The five CC87 subgroups (ST^WU^58-ST^IP^186, ST^WU^58-ST^IP^87-A, ST^WU^155-ST^IP^21, ST^WU^58-ST^IP^87-B, ST^WU^58-ST^IP^24) based on the Warwick University^3^ and Institut Pasteur^2^ MLST schemes are highlighted in color. The host or origin of the strain is highlighted by coloured circles. O-types (*wzm, wzt, wzx* or *wzy*), H-types and fimH-types are represented by coloured rectangles from the inside to the outside. Only antigens found in more than 10 strains are coloured, the others are in grey. The 26 ST58 genomes obtained from the study by Reid et al. are in bold. For the sake of readability, branch lengths are ignored and local support values higher than 0.7 are shown.

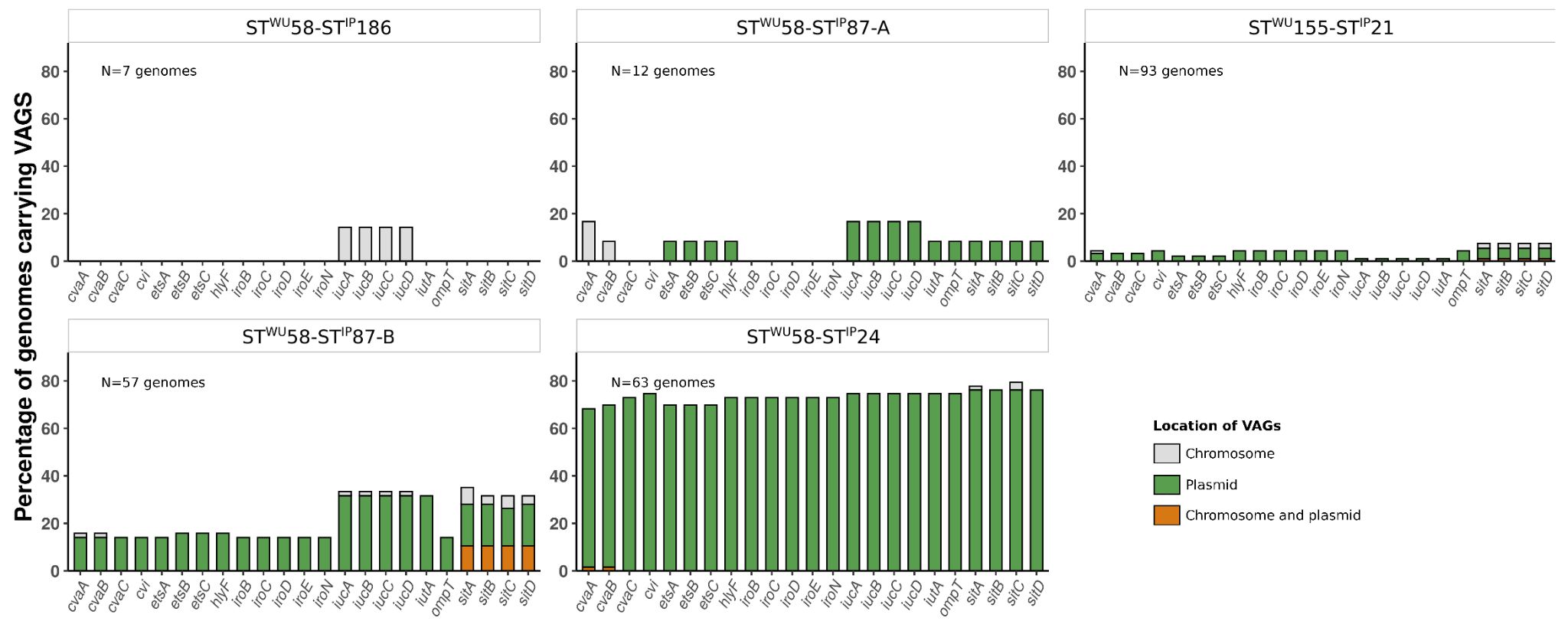

**Figure S3**: Distribution of VAGs and predicted location among the 232 B1 phylogroup CC87 genomes. Results are presented as the percentage of genomes carrying VAGs among each CC87 subgroup depending on their predicted location. Plasmidic location is highlighted in green, chromosomal location in grey and cases with both chromosomal and plasmidic location in orange.

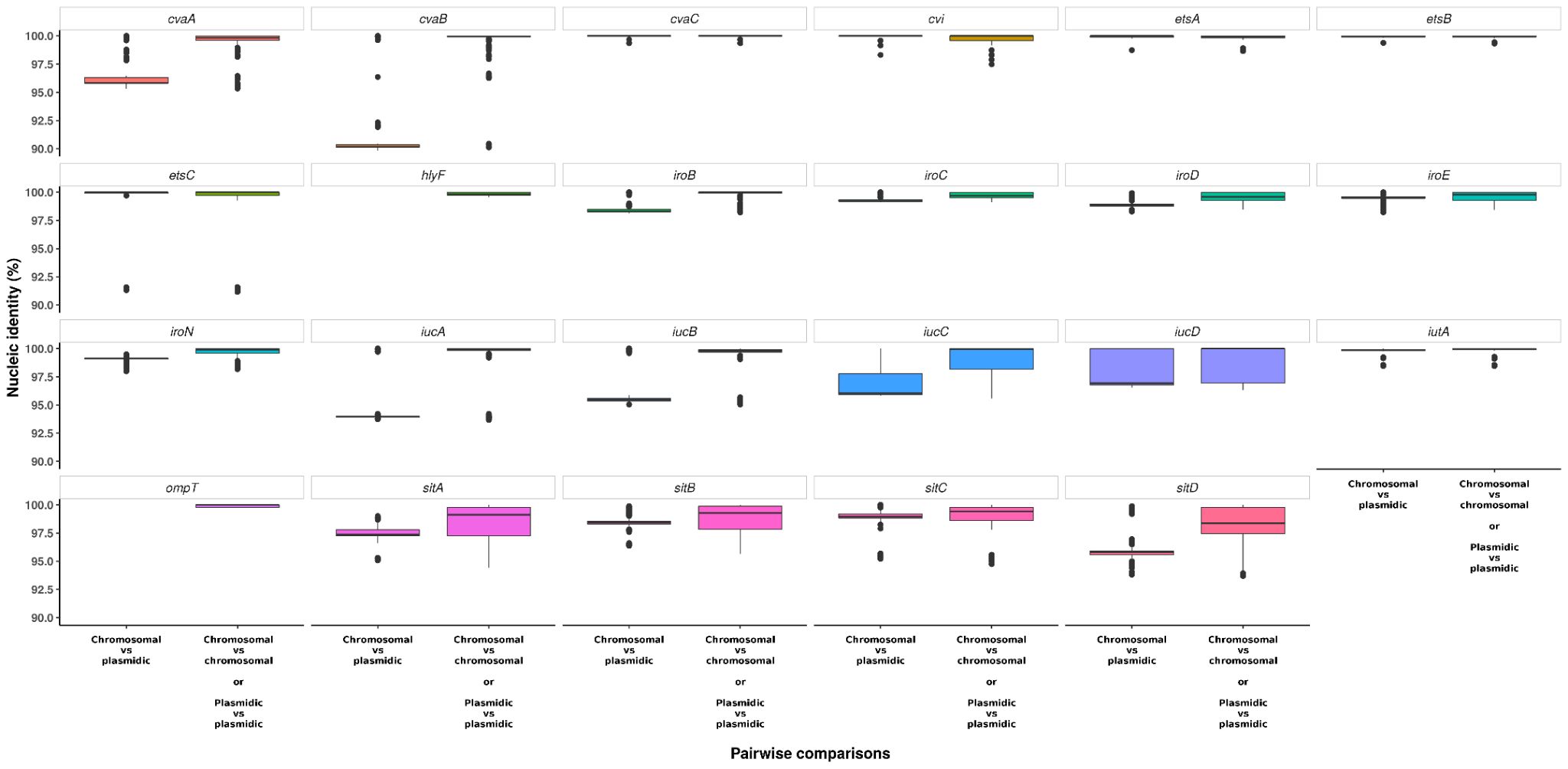
**Figure S4**: Pairwise comparisons of VAG sequences according to their predicted location. We ran an all-against-all blastN analysis among the 370 genomes of *Escherichia*. The nucleic identity distributions of all pairwise comparisons are plotted as boxplots against the predicted location of VAGs within a given pair.

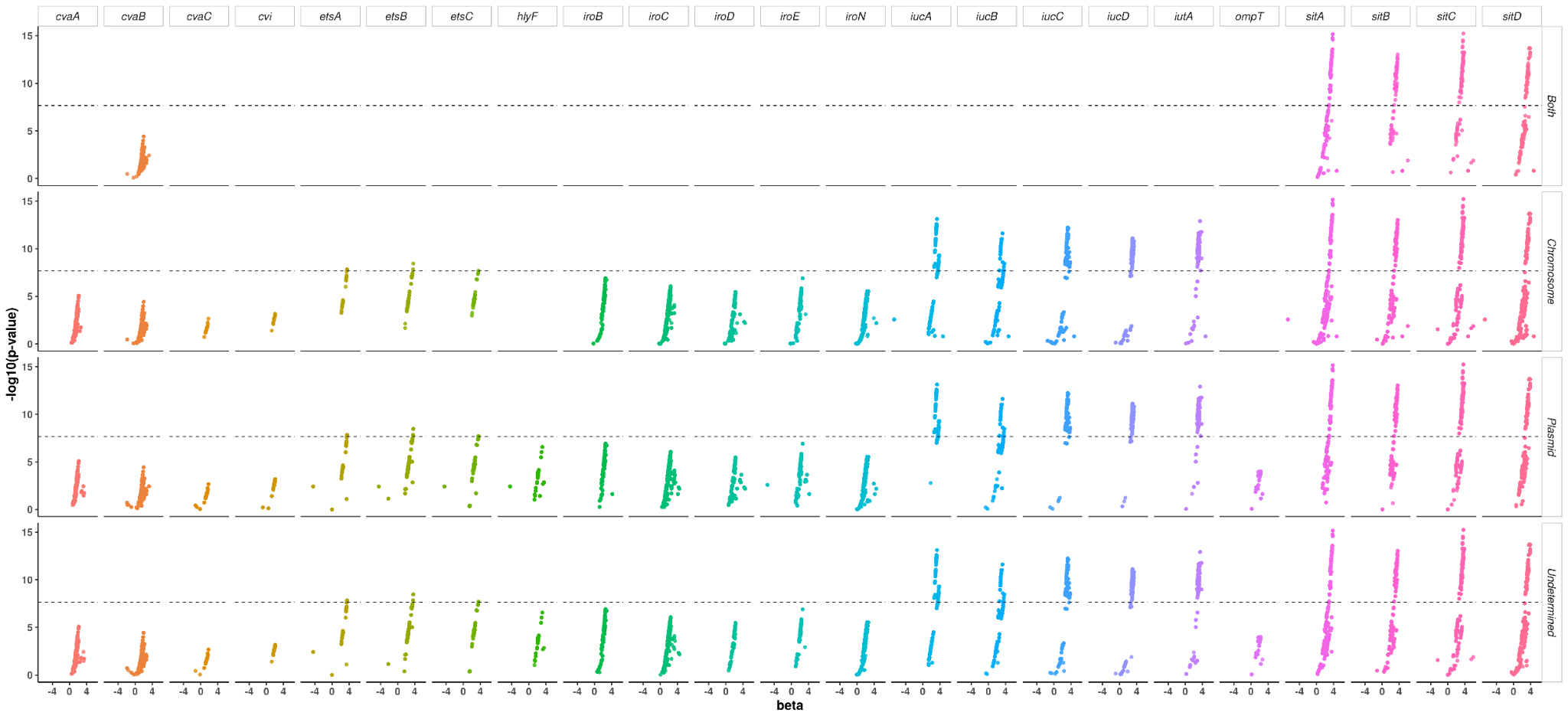

**Figure S5**: Association between virulence and unitigs belonging to the ColV plasmid VAGs at the genus level according to their predicted location. The GWAS results were obtained from a previous study performed on 370 strains of *Escherichia*^4^. The p-value of the association is shown on the y-axis, the effect size (beta) on the x-axis and the significance level of the GWAS analysis with a dotted line. Each facet represents the result for a given VAG in a given predicted location (i.e. chromosomal, plasmidic, both or undetermined).

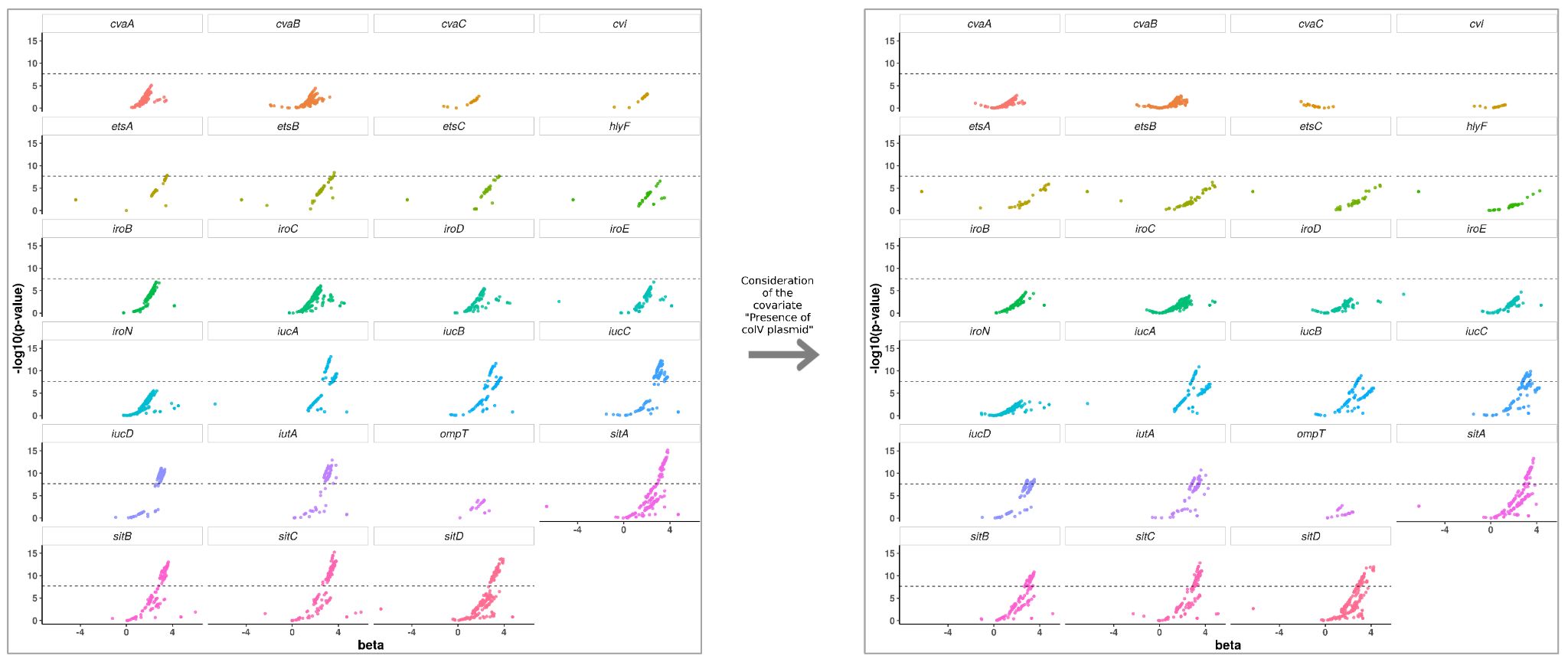

**Figure S6**: Association between the unitigs within each VAG used to infer the presence of the ColV plasmid and virulence in mice after considering ColV plasmids as covariate. The scatter plots on the left hand side of the figure represent the results for the 370 strains of the genus *Escherichia*^4^ without taking into account the presence of the ColV plasmid. The scatter plots on the right hand side of the figure represent the results of the same analysis but considering the presence of the ColV plasmid as a covariate during the GWAS analysis. The p-value of the association is shown on the y-axis, the effect size (beta) on the x-axis and the significance level of the GWAS analysis with a dotted line.

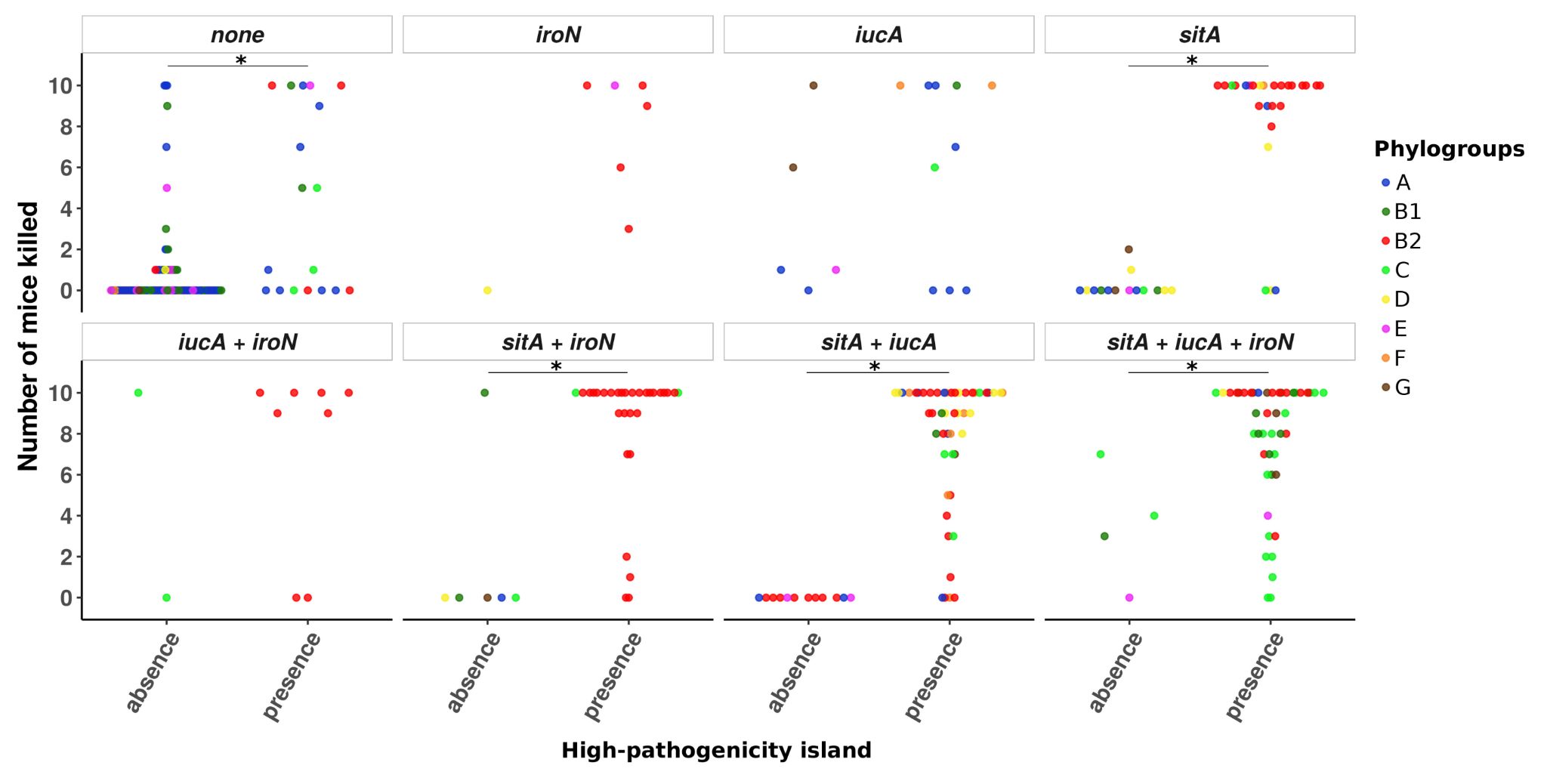

**Figure S7:** Number of mice killed over ten according to the combination of iron acquisition-related VAGs *iroN*, *iucA*, *sitA*. In each facet, each point represents the number of mice killed by a given strain according to the VAG or combination of VAGs it carries. Points are colored according to the phylogroup belonging of the strains.

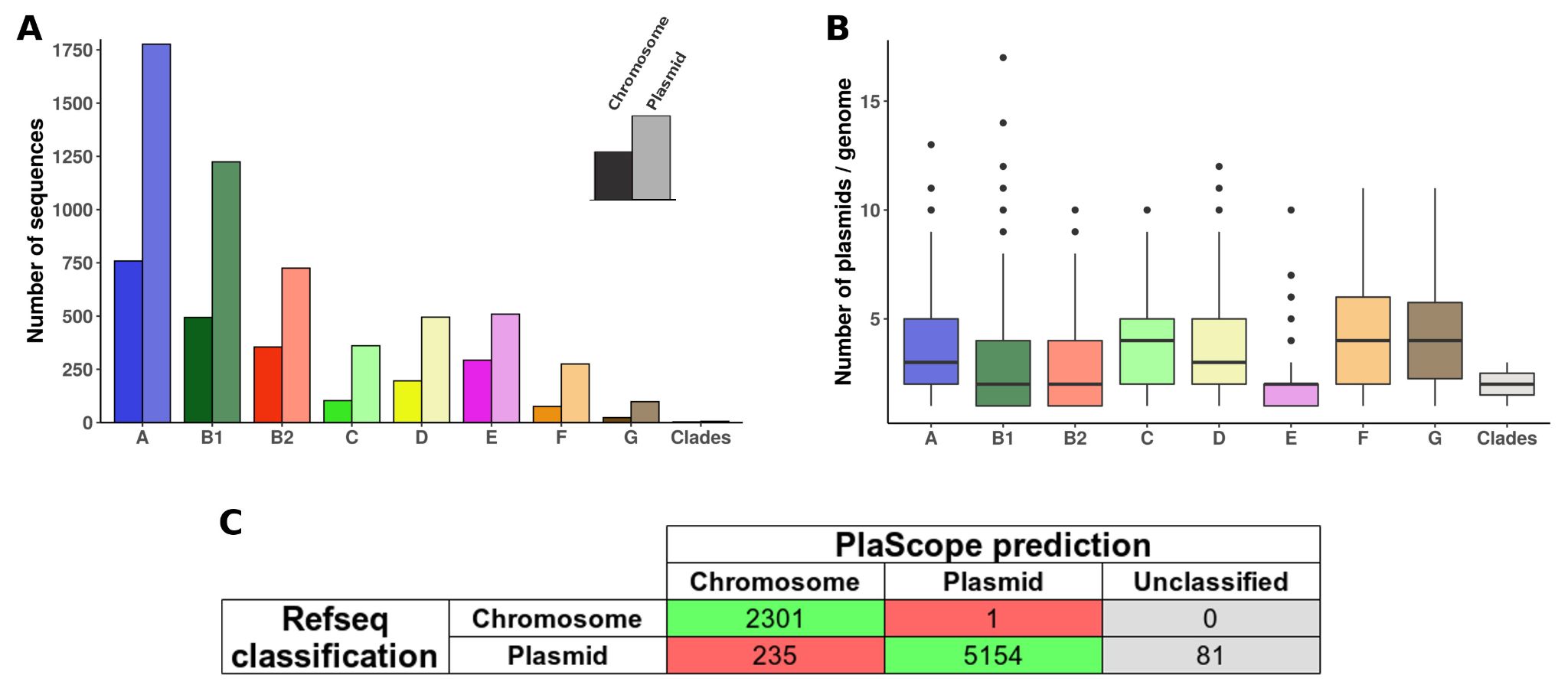
**Figure S8**: Assessment of PlaScope^5^ performances on the 2302 genomes of *E. coli* from RefSeq. A) Total number of chromosomal and plasmidic sequences according to the phylogroup among the RefSeq dataset. B) Distribution of the number of plasmid sequences according to the phylogroup among the RefSeq dataset. C) Comparison of PlaScope and RefSeq classifications. From these results it appears that PlaScope identifies chromosome and plasmid sequences with both high recall (0.94) and specificity (>0.99).

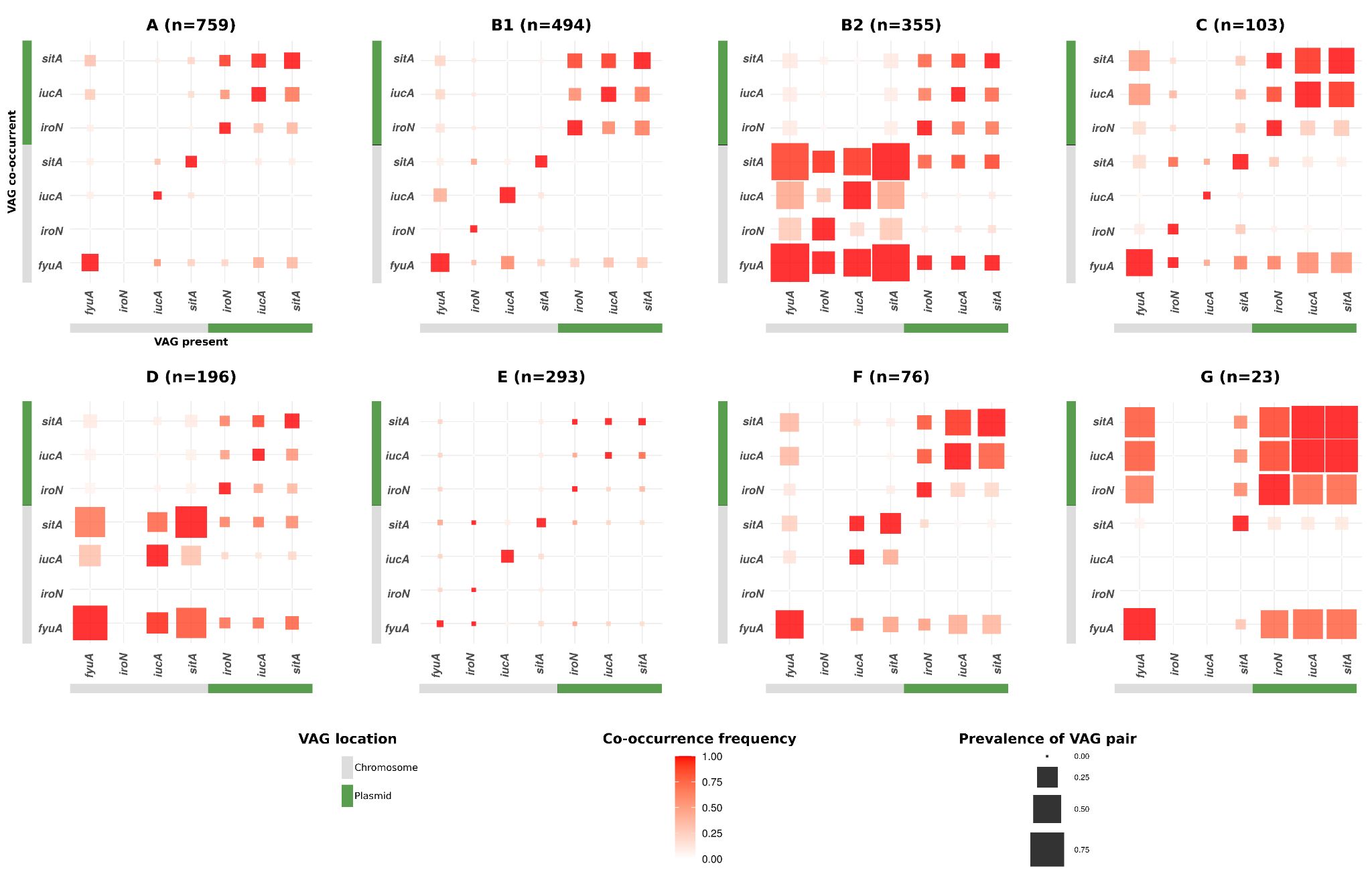

**Figure S9**: Co-occurrence frequency and prevalence of iron acquisition-related VAG pairs among 2299 fully circularized genomes of *E. coli*. For each given VAG on the x-axis, the frequency of co-occurrence with the VAGs on the y-axis is highlighted by a colour gradient. The size of each square is proportional to the prevalence of the VAG pair in the given ST/STc. VAGs are separated according to their location on the chromosome in grey or on the plasmid in green. *Escherichia* Clade I genomes are not included due to their very low prevalence in the data set (n=3).

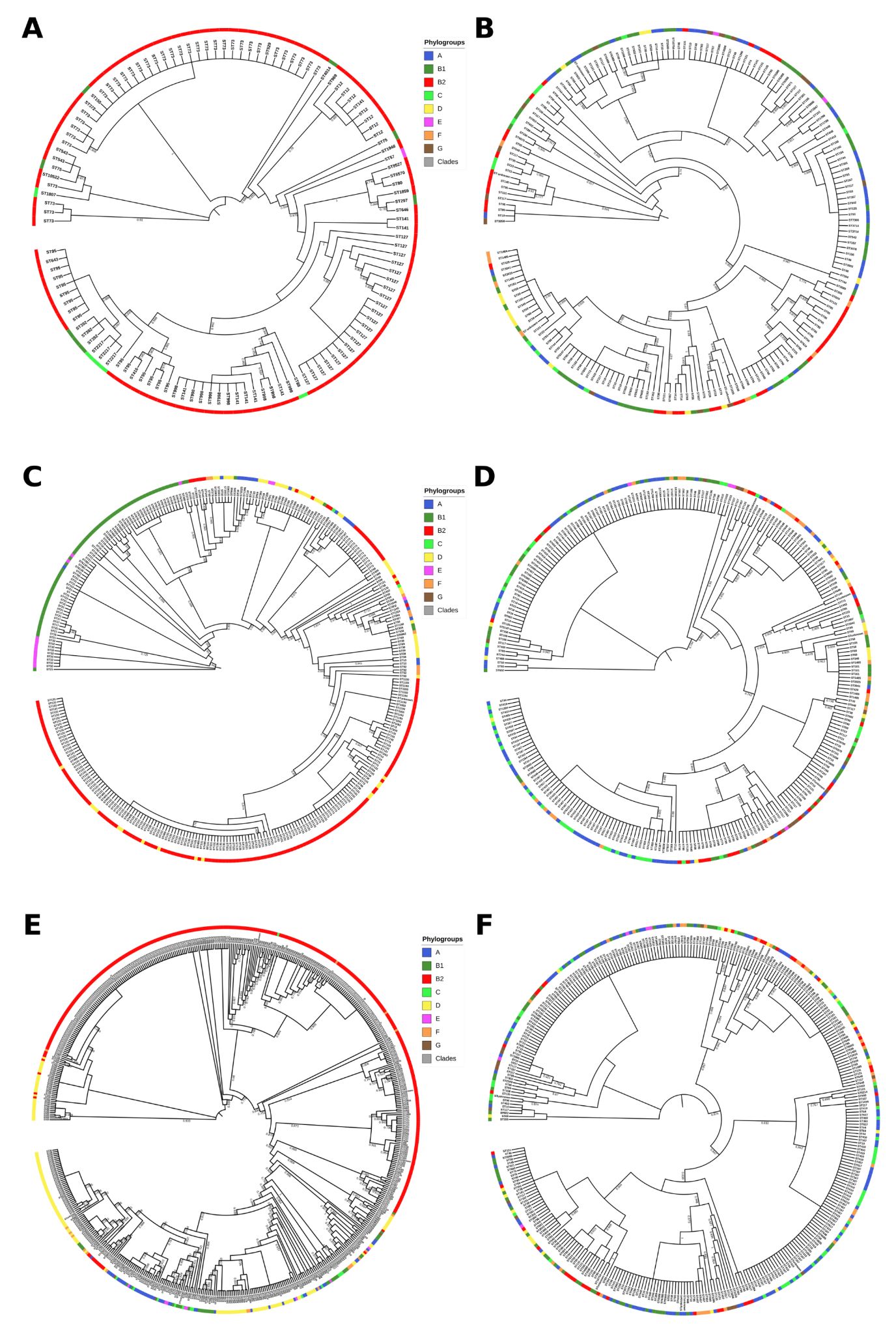

**Figure S10:** Phylogenetic trees of the operon *iro*, *iuc* and *sit*. A) Phylogenetic tree computed from the alignment of chromosomal *iro* operon. B) Phylogenetic tree computed from the alignment of plasmidic *iro* operon. C) Phylogenetic tree computed from the alignment of chromosomal *iuc* operon. D) Phylogenetic tree computed from the alignment of plasmidic *iuc* operon. E) Phylogenetic tree computed from the alignment of chromosomal *sit* operon. F) Phylogenetic tree computed from the alignment of plasmidic *sit* operon. The phylogroup of the strains are highlighted in color in the most outer circles and MLST of the strain is used as leaf label. For the sake of readability, branch lengths are ignored and local support values higher than 0.7 are shown.
